# Celsr3 drives development and connectivity of the acoustic startle hindbrain circuit

**DOI:** 10.1101/2024.03.07.583806

**Authors:** Joy H. Meserve, Maria F. Navarro, Elelbin A. Ortiz, Michael Granato

**Affiliations:** Department of Cell and Developmental Biology, Perelman School of Medicine, University of Pennsylvania, Philadelphia, Pennsylvania, United States of America

## Abstract

In the developing brain, groups of neurons organize into functional circuits that direct diverse behaviors. One such behavior is the evolutionarily conserved acoustic startle response, which in zebrafish is mediated by a well-defined hindbrain circuit. While numerous molecular pathways that guide neurons to their synaptic partners have been identified, it is unclear if and to what extent distinct neuron populations in the startle circuit utilize shared molecular pathways to ensure coordinated development. Here, we show that the planar cell polarity (PCP)-associated atypical cadherins Celsr3 and Celsr2, as well as the Celsr binding partner Frizzled 3a/Fzd3a, are critical for axon guidance of two neuron types that form synapses with each other: the command-like neuron Mauthner cells that drive the acoustic startle escape response, and spiral fiber neurons which provide excitatory input to Mauthner cells. We find that Mauthner axon growth towards synaptic targets is vital for Mauthner survival. We also demonstrate that symmetric spiral fiber input to Mauthner cells is critical for escape direction, which is necessary to respond to directional threats. Moreover, we identify distinct roles for Celsr3 and Celsr2, as Celsr3 is required for startle circuit development while Celsr2 is dispensable, though Celsr2 can partially compensate for loss of Celsr3 in Mauthner cells. This contrasts with facial branchiomotor neuron migration in the hindbrain, which requires Celsr2 while we find that Celsr3 is dispensable. Combined, our data uncover critical and distinct roles for individual PCP components during assembly of the acoustic startle hindbrain circuit.

**Highlights:**

1. The PCP cadherin Celsr3 regulates startle circuit development in zebrafish
2. Celsr3 and other PCP-associated proteins promote Mauthner axon growth and guidance
3. Celsr3 is required for spiral fiber and glia targeting to the Mauthner axon cap
4. Symmetric spiral fiber input to Mauthners is critical for escape direction

## Introduction

The startle response is an evolutionary conserved behavior that enables animals to adopt a protective stance or escape a dangerous situation. In humans, the startle response is elicited by sudden and intense stimuli, which may be auditory, somatosensory, and/or visual, and manifests with an eyeblink and bilateral contraction of facial and neck muscles^1^. In small prey animals including insects^2–4^ and zebrafish^5^, as well as other teleost fish^6^, an intense stimulus can elicit a fast escape. This escape can be directional, away from the perceived stimulus, or in a stereotyped direction, such as forward or backward. The speed and movements of these startle responses depend upon the underlying neural circuit.

In zebrafish, the acoustic startle circuit assembles during embryogenesis^5,7,8^. Central to the circuit are the Mauthner cells, a pair of large reticulospinal neurons located in the hindbrains of some amphibians^9^ and fish^6^, including zebrafish. These neurons are functionally similar to startle-associated giant reticulospinal neurons in the mammalian caudal pontine reticular nucleus^10,11^. In five day old larval zebrafish (5 days post-fertilization, dpf), intense acoustic stimuli perceived by hair cells in the ear and lateral line robustly elicit Mauthner-mediated fast escape responses. Acoustic stimulation of hair cells in the inner ear is conveyed to ipsilateral Mauthners via the eighth cranial nerve (Figure 1A). Mauthner axons cross the midline and extend posteriorly throughout the entire length of the spinal cord, where they synapse onto motor neurons that induce muscle contraction and a sharp turn away from the acoustic stimulus. This initial turn, called the “C1-bend,” is followed by a second turn and subsequent fast swimming to escape (Figure 1B). The circuit also includes feedback inhibitory neurons, which provide both recurrent inhibition (to prevent multiple action potentials from the same Mauthner) and reciprocal inhibition (to prevent both Mauthners from firing)^12^. The Mauthner also receives excitatory input at its axon initial segment from spiral fiber neurons, which are activated in response to acoustic stimuli^13^. These spiral fiber synapses, along with feedback inhibitory synapses and surrounding glia, form the Mauthner axon cap^14^. Unilateral ablations of spiral fiber or Mauthner neurons result in larvae that only escape in a single direction, driven by an initial left C1-bend or right C1-bend^13,15^, which could be towards the source of the acoustic stimulus and danger rather than away. The dire consequences of a misdirected escape demand that proper development of the startle circuit occurs.

**Figure 1:**
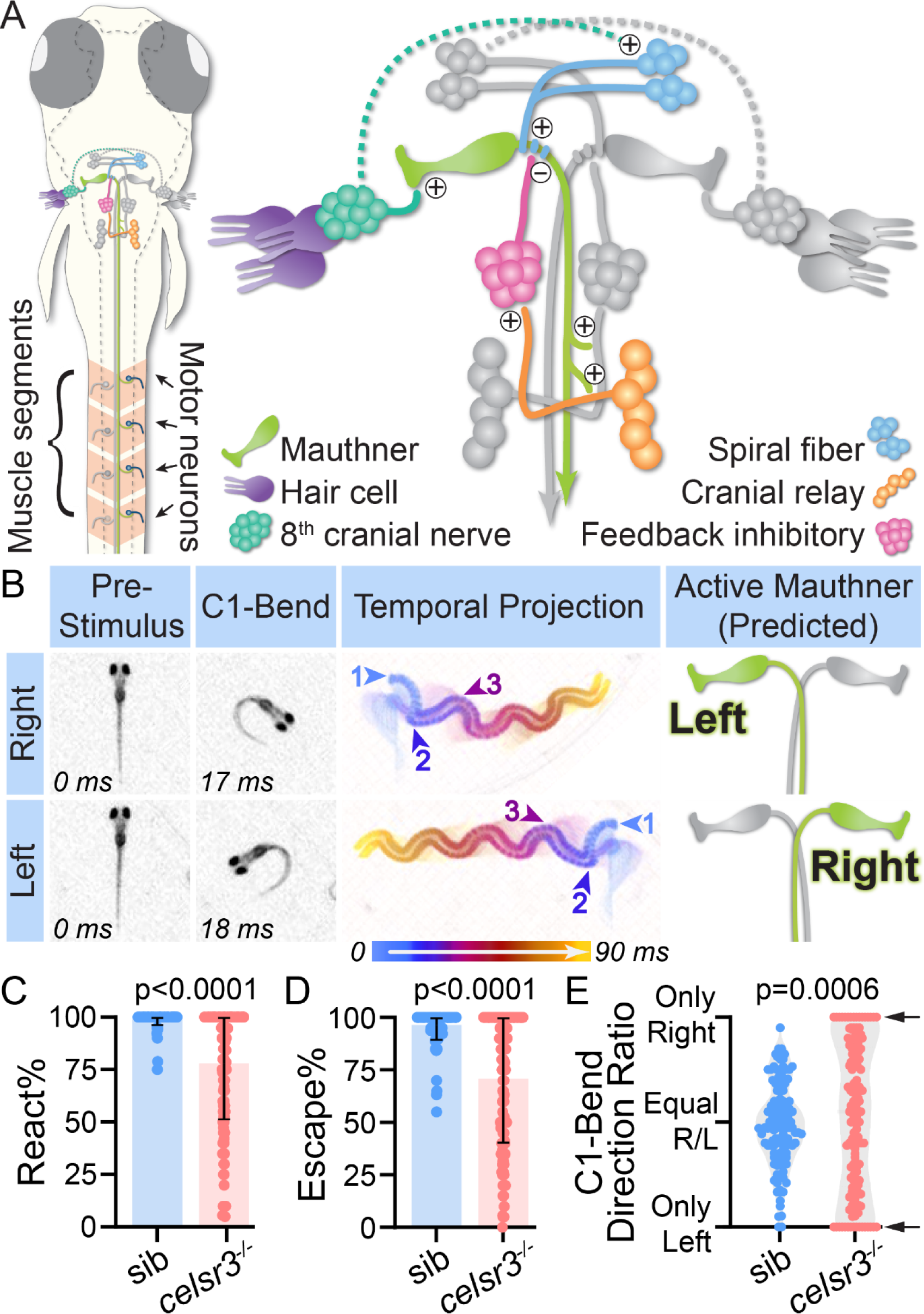
Celsr3 is required for directionally unbiased acoustic startle escapes. A) Schematic of a 5 dpf (days post-fertilization) larva (dorsal view) with a simplified diagram of the hindbrain circuit that drives the acoustic startle fast escape. +/- symbols indicate excitatory/inhibitory synapses. Dotted line indicates unknown connection. In response to a startling acoustic stimulus, Mauthner firing results in direct activation of contralateral motor neurons, leading to unilateral muscle contraction and a sharp turn (C1-bend). B) Representative right and left acoustic startle fast escapes in wild type 5 dpf larvae. Pre-Stimulus and C1-Bend: still images of when the acoustic stimulus was delivered (0 ms) and the highest angle of the Mauthner-dependent initial turn (C1-bend). Middle panel: 90 ms temporal projection of the response, including the C1-bend (begins arrow 1), counter bend (begins arrow 2), and subsequent swimming (begins arrow 3). Active Mauthner: predicted Mauthner that fired to drive right or left escape. C) Percentage of twenty acoustic stimuli that elicited any response (fast escape or response with >17 ms latency, which latency measured as first head movement following stimulus delivery). Each dot represents one fish. n=126 siblings (mix of *celsr3^+/+^* and *celsr3^-/+^*), 154 *celsr3^-/-^* mutants. D) For larvae from (C), percentage of twenty acoustic stimuli that elicited a fast escape (latency ≤17 ms). 47% of mutant larvae responded within the sibling range (88-100%). E) For larvae from (D) that performed at least six fast escapes, initial C1-bend direction ratio (right versus left) for individual larva. Arrows indicate larvae that only turn one direction (“100% biased”; 24% of mutants). n=126 siblings, 133 mutants.

Neural circuit formation requires coordination of numerous developmental steps. Neurons must properly differentiate, migrate, if necessary, to their correct location, extend axons, and form synapses with appropriate targets. Directing all these processes are molecular pathways and proteins. Here, we focus on the planar cell polarity (PCP) pathway, which plays numerous roles in nervous system development. These roles include tissue polarization during neural tube closure^16–18^, polarization of interacting cells during neuron migration^19–22^, axon guidance^23,24^, dendrite development^25–27^, and synapse formation^28^. These processes do not necessarily require all PCP core proteins or follow the classic epithelial patterning model of protein interactions^29^. Celsrs (Cadherin EGF LAG seven-pass G-type receptors) are core PCP pathway components homologous to *Drosophila* Flamingo/Starry night^30–32^. Mammals have three *CELSR* genes (*CELSR1*, *CELSR2*, and *CELSR3*)^33^, while zebrafish have four (*celsr1a*, *celsr1b*, *celsr2*, and *celsr3*)^34^. Results from numerous studies suggest CELSR1 primarily regulates epithelial PCP, while CELSR2 and CELSR3 have both overlapping and unique roles in neural development^32,35^. The precise roles different Celsrs play in neural development and whether other PCP-pathway proteins are required are complex questions not yet fully understood.

Our interest in the roles of PCP proteins in startle circuit development began with a fortuitous observation: while studying the role of *celsr3* in axon regeneration^36^, we discovered uninjured *celsr3* mutant larvae display a startle behavior deficit. In response to non-directional acoustic stimuli, wild type larvae initiate left escapes and right escapes with roughly equal probabilities (“unbiased escapes”). In contrast, we find that *celsr3* mutant larvae have a significantly higher probability of executing startle escapes in only one direction, left or right (“biased escapes”). We identify the underlying cause of this behavioral deficit in *celsr3* mutants by demonstrating Celsr3 is required for axon growth and guidance of both Mauthner cells and spiral fiber neurons, and disruption of either neuron type can lead to biased escapes. We additionally set out to determine the functional requirements of Celsr2 and Frizzled 3a (Fzd3a) in startle circuit development, as distinct Celsrs can have overlapping functions, and Celsr proteins frequently act in conjunction with Frizzled PCP proteins. While loss of *celsr2* or *fzd3a* alone do not disrupt startle circuit development or behavior, double mutants of *celsr3* and *celsr2* or *fzd3a* display strong defects, suggesting redundancy within PCP-pathway proteins that ensures proper startle circuit development. By analyzing Mauthner axonal development in *celsr3^-/-^;celsr2^-/-^* embryos, we find that Mauthner axons frequently grow anteriorly instead of posteriorly, and Mauthner cells are absent by the end of embryogenesis, presumably due to cell death. Our data also reveals diverging roles for *celsr3* and *celsr2* in hindbrain development. *celsr2* plays a well-documented critical role in facial branchiomotor migration^19,21,37^ while we find that *celsr3* is dispensable for this process. Finally, we utilize Fzd3a dominant-negative transgenic spatiotemporal expression to demonstrate that guidance to the axon cap for spiral fiber axons glia guidance is at least partially independent from the presence of Mauthner cells, suggesting additional unknown cues guide formation of the axon cap. Combined, our results reveal partially overlapping and unique roles for Celsrs in hindbrain development, with Celsr3 directing assembly of pre- and post-synaptic partners within the acoustic startle circuit.

## Results

### Celsr3 is required for unbiased acoustic startle escapes

Behavioral output is a powerful indicator of circuit function and integrity. This is especially true for the acoustic startle escape response in zebrafish (Figure 1A,B), as neurons comprising the underlying circuit have been well-characterized, and the behavioral phenotypes following manipulation (ablation, silencing, and/or activation) of many of these neurons are known^13,15,38–42^. In response to intense and sudden acoustic stimuli, 5 dpf larvae have a high reaction frequency (Figure 1C) and primarily respond with fast escapes (Figure 1D). Directional acoustic stimuli elicit escapes away from the startling stimulus, driven by the neural circuitry underlying the response (Figure 1A,B). Following a non-directional acoustic stimulus, larvae perform an escape maneuver to the left or to the right, following the initial Mauthner-dependent C1-bend direction, without bias. Over multiple stimuli, wild type and *celsr3* sibling larvae on average perform equal numbers of escapes to the left and to the right (measured as ratio of right C1-bends versus left C1-bends for individual larvae; Figure 1E). In contrast, the C1-bend direction ratio is unbalanced in *celsr3* mutant larvae, with 29% only initiating escapes in one direction over multiple stimuli (Figure 1E). Additionally, *celsr3* mutants on average exhibit a reduction in total acoustic response frequency (includes fast escapes and slower responses; Figure 1C) and fast escape frequency (Figure 1D), though 47% of *celsr3* mutant larvae react at sibling levels (within one standard deviation from sibling average, i.e. 88-100% fast escape frequency) (Figure 1D). The reduction in response frequency may be caused by defects outside the startle circuit, as *celsr3* mutants lack a swim bladder, exhibit reduced spontaneous movements (sibling bouts/min average=72, n=27; mutant bouts/min average=31, n=21; p-value<0.001), and are not viable past ∼11 dpf (*celsr3^fh^*^339^ is a presumed null allele^36^). Nonetheless, the biased escapes we observe in *celsr3* mutants suggest *celsr3* has a specific role in startle circuit development and/or function.

### Celsr3 is required for Mauthner cell development

We first assessed Celsr3’s role in Mauthner cell development, since the initial startle escape turn direction is dependent on whether the left or right Mauthner cell fires (Figure 1A,B). Given that *celsr3* mutants were more likely to perform escapes in one direction, we hypothesized that *celsr3* mutants may have deficits in Mauthner cell development and/or connectivity. To visualize Mauthner cells, we utilized the *Tol-056* transgenic line^43^, which expresses cytosolic GFP in Mauthners, and the neurofilament antibody 3A10, which labels Mauthner axons. At 5 dpf, 100% of wild type and *celsr3* sibling larvae have two Mauthner cells (Figure 2A,C), while ∼10% of *celsr3* mutant larvae lack one Mauthner cell (Figure 2B,C). We do not observe any severe morphological defects in Mauthner somas or hindbrain axons in *celsr3* mutants with one or two Mauthner cells (Figure 2B). We next assessed Mauthner axons in the spinal cord in *celsr3* mutants. During normal development, each Mauthner projects a single axon that crosses the midline in the hindbrain and extends posteriorly into the spinal cord (Figure 2D). By 5 dpf, the Mauthner axon measures ∼2700μM in total length and extends along the entire spinal cord, which is surrounded by repeating muscle segments. We used these segments as landmarks to quantify Mauthner axon length. At 5 dpf, Mauthner axons in sibling larvae extend at least to segment 30 (Figure 2D,F,H). In contrast, Mauthner axons in *celsr3* mutants are significantly shorter, as 50% of axons do not reach segment 30 (Figure 2E,G,H), and some were as short as segment 15 (Fig. 2H). These results demonstrate *celsr3* is critical for proper Mauthner development and axon growth.

**Figure 2:**
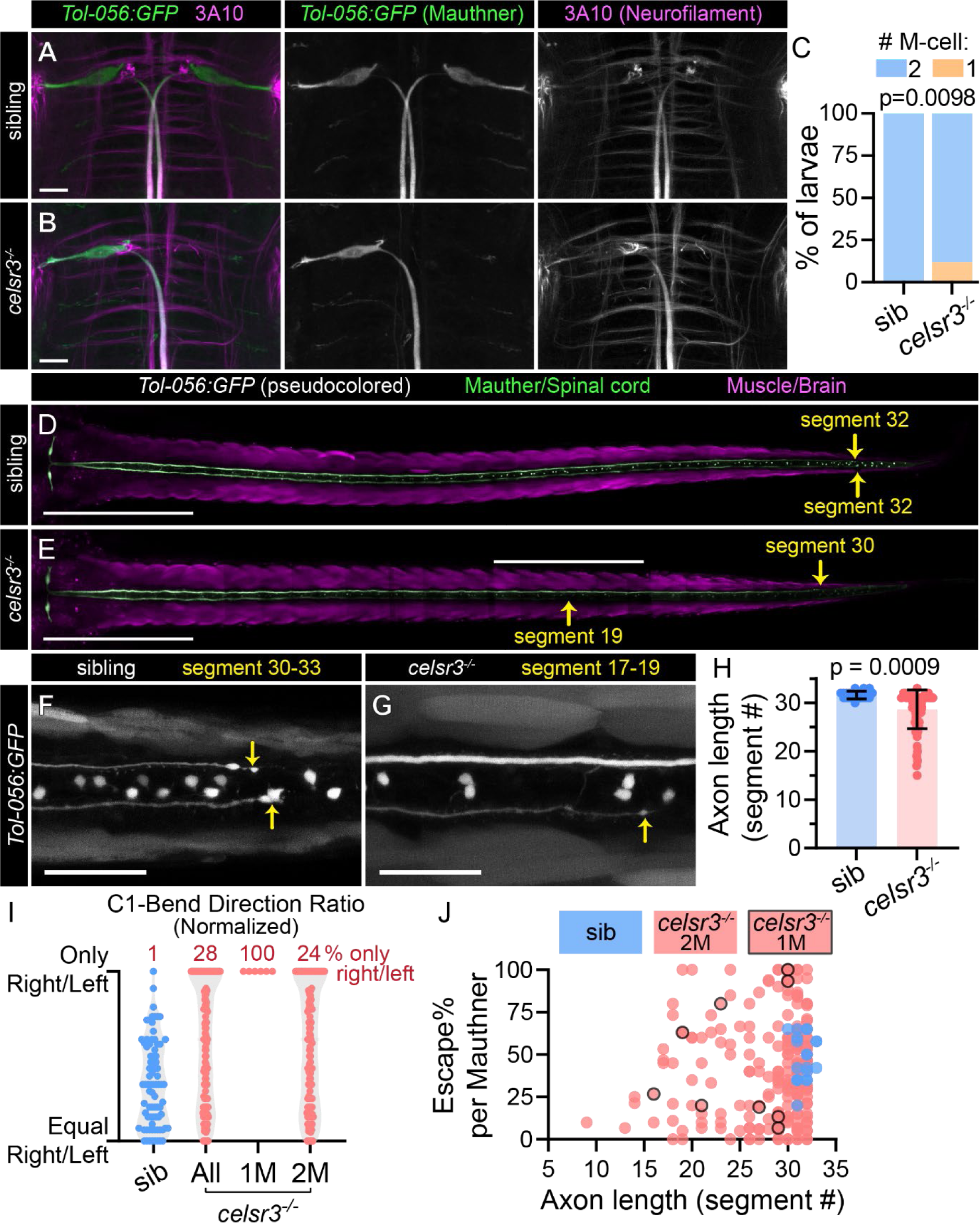
Celsr3 is required for Mauthner development. A,B) *Tol-056* transgene expression of cytosolic GFP in Mauthner neurons in *celsr3* sibling (A) and mutant (B) larvae. Mauthner axons are labeled with α-3A10 (neurofilament antibody) staining. All images and quantifications in figure are of 5 dpf larvae. Scale bar=20μM, Z-projection. C) Mauthner cell number in *celsr3* siblings (100% 2 M-cells; n=51) and *celsr3* mutants (12% 1 M-cell, 88% 2 M-cells; n=67). D,E) Mauthner somas and axons in *Tol-056 celsr3* sibling (D) and mutant (E). The Mauthner/spinal cord is pseudo-colored green for ease of visualization, while non-Mauthner/spinal cord GFP expression is magenta. Only z-frames with the Mauthner axon in focus are included. Arrows indicate where the Mauthner axons end and corresponding muscle segments, numbered anterior to posterior. Scale bar=500μM, Z-projection. F,G) Magnified Mauthner axon ends in *Tol-056 celsr3* sibling (F) and mutant (G). The top axon in (G) continues to segment 30. Scale bar=50 μM, Z-projection. H) Mauthner axon lengths in *celsr3* siblings and mutants measured by muscle segment containing the Mauthner axon end. Average sibling=segment 32 (n=22 axons), mutant=segment 29 (n=66 axons). I) Escape C1-bend ratio for *celsr3* siblings and mutants that performed at least six fast escapes in response to 20 acoustic stimuli, normalized so all fish turning only one direction, left or right, appear at the same y position. n=79 siblings, 6 mutants with one Mauthner (1M), 109 mutants with two Mauthners (2M). J) Escape frequency (Escape%) for individual Mauthners was calculated using total Escape% and turn bias ratio for individual fish; left C1-bend escape was assigned to right Mauthner, right C1-bend escape assigned to left Mauthner. n=20 sibling Mauthners, 253 *celsr3* mutant Mauthners. Points outlined in black indicate *celsr3* mutant Mauthners from larvae with a single Mauthner (1M). Individual Mauthner escape% and axon length are not significantly correlated (simple linear regression, p=0.63) in *celsr3* mutants.

We next asked whether Mauthner defects in *celsr3* mutants are correlated with biased escapes. Similar to previous work in larvae with only a single Mauthner following ablation of the second Mauthner^13,15^, *celsr3* mutant larvae with a single Mauthner cell initiate escapes to only one side, contralateral to the remaining Mauthner (Figure 2I). Unexpectedly, *celsr3* mutant larvae with two Mauthner cells also exhibit biased escapes (Figure 2I). To explore the possibility that Mauthner axon length might relate to Mauthner activity, we calculated escape frequency for individual Mauthners. We used the escape frequency for an individual larva and that larva’s turn bias to calculate response frequency for individual Mauthners, with the assumption that left escapes were driven by right Mauthner firing, and vice versa. Mauthner escape frequency and Mauthner axon length do not correlate, and there is no clear correlation between mutants with single Mauthners and Mauthner axon length or escape% (Figure 2J). We conclude that shortened Mauthner axons do not affect escape frequency or direction bias in *celsr3* mutants. In contrast, loss of Mauthner cells does contribute to biased escapes, though 81% of *celsr3* mutants with completely biased escapes have two Mauthners. Therefore, we predicted that *celsr3* is required in additional startle circuit neurons critical for unbiased escapes.

### Celsr3 is required for spiral fiber axon guidance

Mauthner cells receive excitatory input from the eighth cranial nerve and from spiral fiber neurons that are activated by acoustic stimuli^13^. Spiral fibers form mixed electrical and glutamatergic synapses on the Mauthner axon initial segment (AIS)^14,44,45^. Excitatory spiral fiber synapses and competing feedback inhibitory synapses at the Mauthner AIS are surrounded by astrocyte-like glial cells, which all together comprise the Mauthner axon cap^46^ (Figure 3A-J). To visualize spiral fiber somas and axons, we utilized the transgenic reporters *hcrt:Gal4, UAS:Kaede* (*hcrt::Kaede*)^13^, which stochastically labels spiral fiber neurons (Figure 3D,I), and *j1229a*^47^, which expresses cytosolic GFP in various neuron populations, including spiral fiber and Mauthner neurons. In wild type larvae, approximately ten spiral fiber soma per brain hemisphere are organized in two bilateral groups rostroventral to the Mauthner cells^45^ (Figure 3A-D). These spiral fiber neurons project axons across the midline, then turn posteriorly to establish synapses at the Mauthner AIS (Figure 3A,B,D). At the axon cap (Figure 3F-J), a nuclei-free region (Figure 3G) is surrounded by S100B+^48^ glia (Figure 3H) and contains spiral fiber axons (Figure 3I) wrapping around the Mauthner AIS (Figure 3J). In *celsr3* mutants, while some spiral fiber axons project correctly to the Mauthner AIS, we also observe misprojected spiral fiber axons that form ectopic bundles reminiscent of those at the endogenous axon cap (Figure 3K,L). To further characterize spiral fiber defects in *celsr3* mutants, we utilized an antibody against the potassium channel subunit Kv1.1, which is highly expressed in spiral fiber axons^49^, particularly at the axon cap (Figure 3M). We find that Kv1.1 localizes to the axon cap in all *celsr3* mutants (n=48), suggesting at least some spiral fibers axons have reached the Mauthner (Figure 3N). In addition to this wild-type-like innervation pattern, we also observe ectopic accumulations of Kv1.1 rostroventral to the Mauthner AIS along the path of spiral fiber projections (Figure 3N), indicative of misprojecting spiral fiber axons. These ectopic accumulations, which we term “ectopic caps” for their resemblance to spiral fiber axons and Kv1.1 at endogenous Mauthner axon caps, are present in *celsr3* mutant brain hemispheres with or without a corresponding Mauthner cell (Figure 3O,P), most commonly at the position of spiral fiber somas along the anterior-posterior axis (Figure 3Q,R). These ectopic caps also co-localize with the gap junction protein Connexin 35/Cx35 (Figure 4B-D) which marks spiral fiber synapses with the Mauthner^50^, indicating misguided axons form synapses with inappropriate targets. These data indicate *celsr3* is required for spiral fiber axon guidance to the Mauthner AIS.

**Figure 3:**
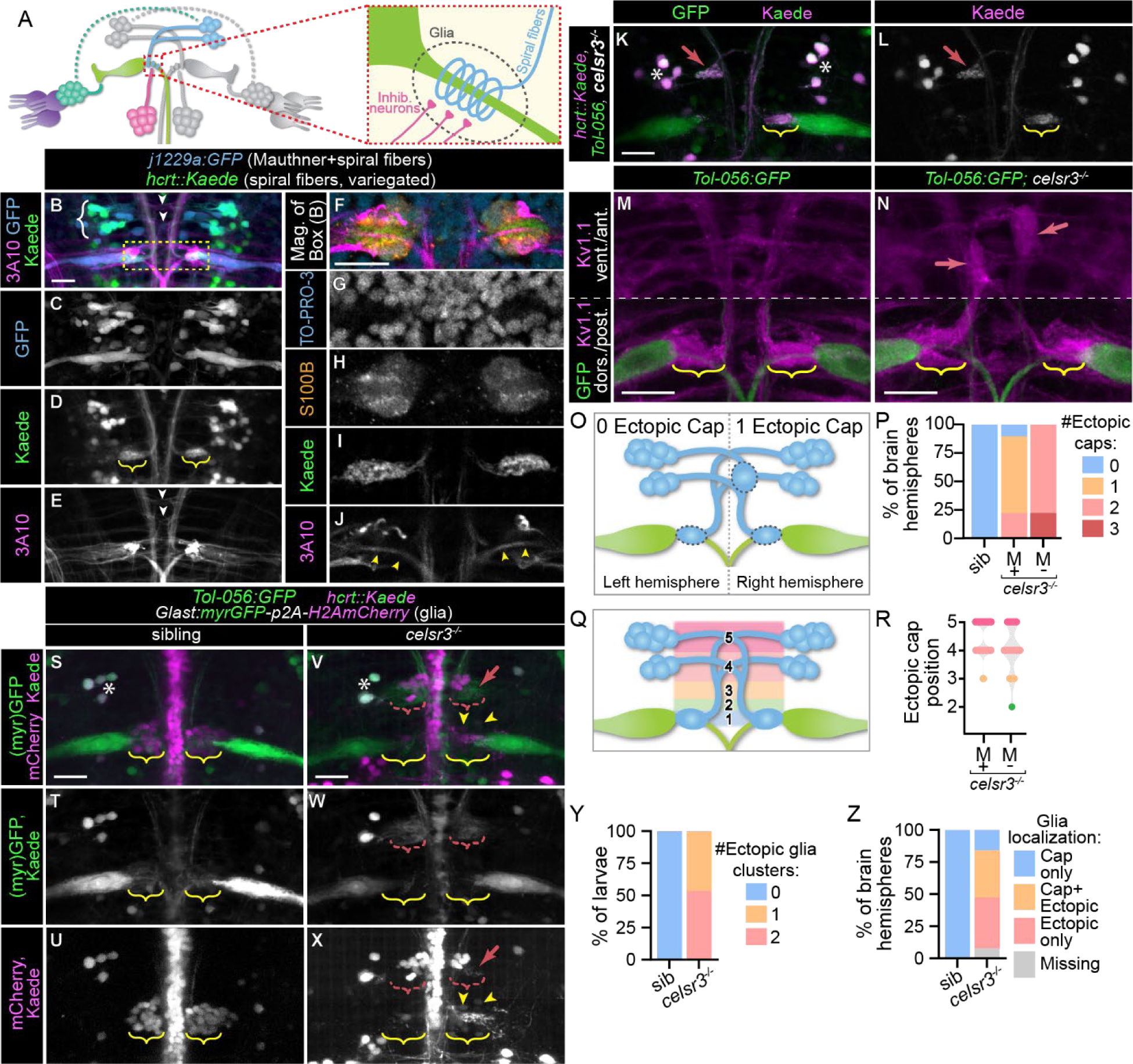
Celsr3 is required for spiral fiber axon guidance and glial localization to the axon cap. A) The axon cap (dotted red box in circuit diagram) is comprised of excitatory spiral fiber axons (blue) and feedback inhibitory axons (pink) synapsing with the Mauthner (green) axon initial segment (AIS), surrounded by astrocyte-like glia (dotted line). B-E) A wild type larva with *j1229a* transgene expression of cytosolic GFP in Mauthner and spiral fiber neurons (B,C), *hcrt:Gal4, UAS:Kaede/hcrt::Kaede* transgene expression in a subset of spiral fibers (B,D; protein mostly converted from green to red), and α-3A10 staining (B,E). White brackets in (B) indicate spiral fiber somas. Yellow brackets in (D) indicate spiral fiber axons at the axon cap. White arrows in (E) indicate spiral fiber axons at midline crossing. All images and quantifications in figure are of 5 dpf larvae. Scale bar=20μM, Z-projection. F-J) Magnification of box in (B) showing both Mauthner axon caps with TO-PRO-3 labeling nuclei (F,G), α-S100B labeling glia (F,H), Kaede labeling spiral fiber axons (F,I), and α-3A10 labeling Mauthner axons (F,J). Yellow arrowheads in (J) indicate Mauthner AISs. Scale bar=20μM, single Z-frame. K,L) *celsr3* mutant larva with *Tol-056:GFP* (K) and *hcrt::Kaede* (K,L) transgene expression. White asterisks indicate spiral fiber soma (note: only a subset of spiral fiber neurons are labeled due to variegated transgene expression). Yellow brackets indicate normal spiral fiber axon targeting to Mauthner axon cap. Red arrow indicates inappropriate spiral fiber axon targeting. Scale bar=20μM, Z-projection. M,N) Mauthner axon cap region of sibling (M) and *celsr3* mutant (N) *Tol-056:GFP* larvae. Kv1.1 localizes to Mauthner axon caps (yellow brackets) that spiral fiber axons target and accumulates at ectopic positions (red arrows) where spiral fiber axons frequently misproject. Scale bar=20μM, Z-projection (dorsal range included for posterior Mauthner axon cap region and ventral range included for anterior spiral fiber axon misprojections, for clarity). O) Schematic for quantifying spiral fiber misprojections, measured as accumulation of Kv1.1 in ectopic “axon caps,” as in (N) (red arrows), within 60μM anterior to Mauthner AIS and 60μM lateral to midline (boundaries of spiral fiber soma). Left brain hemisphere is normal, with 0 ectopic caps, while right brain hemisphere has 1 ectopic spiral fiber cluster. For quantification in (P), this example would be counted as 0 (for left) and 1 (for right). P) Quantification of “ectopic caps,” presumed to represent spiral fiber misprojections, assessed by counting numbers of large Kv1.1 accumulation (≥5μM diameter) rostroventral to position of normal axon cap, as in (O). For each larva, the left and right hindbrain were counted independently (two “hemispheres” per larva). In addition to ectopic Kv1 accumulation, all sibling and *celsr3* mutant hemispheres with a Mauthner present had Kv1.1 accumulation at the endogenous axon cap region surrounding the Mauthner AIS. *celsr3* mutant brain hemispheres where Mauthner neurons were absent only had ectopic caps. n=10 sibling hemispheres, 58 *celsr3* mutant hemispheres with Mauthners, 9 *celsr3* mutant hemispheres without Mauthners. Q) Schematic for quantifying anterior-posterior position of ectopic caps quantified in (P). 1= positioned at Mauthner AIS, 2=<20μM from correct position, 3=posterior to spiral fiber soma, 4=at posterior spiral fiber midline crossing, 5=at anterior spiral fiber midline crossing. For example, ectopic cap in (O) would be counted as 4. When the Mauthner neuron was absent, additional axon tracts labeled by Kv1.1 were used to identify where correct axon cap positioning was expected. R) Quantification of anterior-posterior position of ectopic caps from (P). For brain hemispheres with two ectopic caps, each position was counted separately. n=63 *celsr3* mutant ectopic caps in hemispheres with Mauthners, 17 *celsr3* mutant ectopic caps in hemispheres without Mauthners. S-X) Sibling (S-U) and *celsr3* mutant (V-X) larvae with *Glast:myrGFP-p2A-H2AmCherry* (membrane GFP, nuclear mCherry) transgene labeling astrocytes, as well as *Tol-056:GFP* and *hcrt::Kaede* (partially converted from green to red) transgenes. Green channel (myrGFP, GFP, and Kaede) in S,V,T,W; red channel (H2AmCherry, Kaede) in S,V,U,X. White asterisks indicate spiral fiber soma (note: only a subset of spiral fiber neurons are labeled due to variegated transgene expression). Yellow brackets indicate endogenous Mauthner axon cap, red brackets indicate ectopic localization of glia, red arrow indicates misprojected spiral fiber axons, and yellow arrowheads indicate two example glia localized correctly to the endogenous axon cap. Scale bar=20μM, Z-projection. Y) Quantification of percent of sibling or *celsr3* mutant larvae with zero, one (left or right), or two (left and right) anterior ectopic glial clusters. All larvae for quantification had two Mauthners. n=16 siblings, 15 *celsr3* mutants. Z) For larvae in (Y), glia on one side of the hindbrain (left or right hemisphere) were categorized as correctly localized to only the endogenous axon cap at the Mauthner AIS, localized to both the endogenous axon cap and an anterior ectopic location, localized only to an ectopic location, or not observed/missing. n=32 sibling hemispheres, 30 *celsr3* mutant hemispheres.

**Figure 4:**
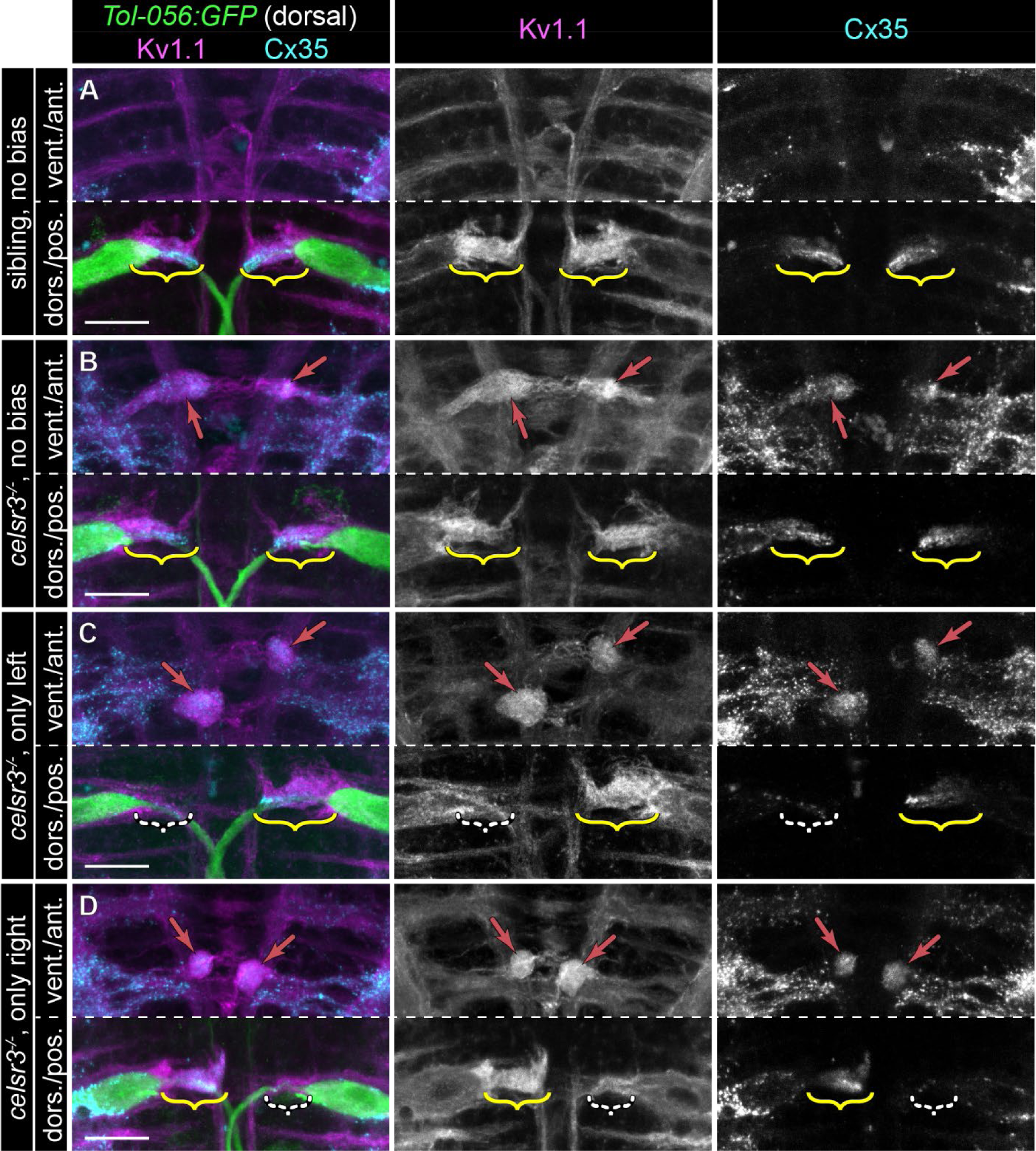
Asymmetric spiral fiber input to Mauthner neurons in *celsr3* mutants correlates with biased escapes. A-D) Region of hindbrain with Mauthner and spiral fiber neurons from 5 dpf *Tol-056:GFP* larvae in (A) siblings with unbiased escapes (40-60% right/left C1-bend direction ratio; all categories ≥6 escapes over 20 stimuli), (B) *celsr3* mutants with unbiased escapes, (C) *celsr3* mutants that only made left C1-bends (right Mauthner predicted active), and (D) *celsr3* mutants that only made right C1-bends (left Mauthner predicted active). Composite images are Z-projections with ventral range for anterior portion and dorsal range for posterior portion. Kv1.1 (left and middle panels) localizes to endogenous axon caps (brackets) and ectopic caps (red arrows in B-D) presumed to represent spiral fiber misprojections. α-Cx35 (left and right panels) labels spiral fiber synapses at endogenous (A-D) and ectopic (B-D) caps. White dashed brackets in (C,D) indicate the axon cap with reduced Kv1.1/Cx35 signal, which corresponds to the inactive Mauthner, based on escape bias. Scale bar=20 μM.

Previous work has shown that astrocyte-like glia surrounding spiral fiber and inhibitory synapses arrive at the axon cap after spiral fiber axons appear^14,51^, yet the signals guiding them to this location are unknown. We utilized a transgenic line labeling astrocytes, *Glast:myrGFP-p2A-H2AmCherry*^52^, to visualize axon cap glia (Figure 3S-X).

In *celsr3* mutants, glia co-localize with spiral fiber axons at the bonafide Mauthner axon cap but also localize to rostroventral positions and with misprojected spiral fiber axons (Figure 3V-Z). The glia mislocalization phenotype is variable in *celsr3* mutants, similar to misguided spiral fiber axons, and we observed at least one ectopic glial cluster in the region of the Mauthner and spiral fiber somas in all *celsr3* mutants imaged (Fig. 3Y). While we always observe Kv1.1 localization to the Mauthner axon cap in *celsr3* mutants, we failed to detect *Glast+* glia at the axon cap in ∼45% of *celsr3* mutant brain hemispheres with Mauthner cells (Figure 3Z). This suggests spiral fiber axons may play a role in attracting glia to the axon cap, such that misguided spiral fiber axons can redirect glia away from their proper position. Thus, *celsr3* is required for glial guidance to the axon cap, either indirectly by promoting spiral fiber axon growth and guidance or directly by functioning in axon cap glia guidance.

### Spiral fiber input at the axon cap is correlated with startle turn bias

Spiral fiber input is required for a robust, fast startle escape^13^. Larvae with unilaterally ablated spiral fiber soma only execute fast escapes in one direction^13^, similar to the defects observed in *celsr3* mutants. We wondered if stochastic defective excitatory input to the left and right Mauthners correlates with and might be causative of biased escapes. In *celsr3* siblings, Kv1.1 and Cx35 localize to both the left and right Mauthner axon caps (Figure 4A). Similarly, in mutants with unbiased C1-bend direction ratio (40-60% right escapes), Kv1.1 and Cx35 are symmetrically localized to the left and right axon caps (n=9/12 larvae with symmetrical Kv1.1 at caps; Figure 4B). In contrast, in *celsr3* mutants that are 100% left-biased, indicating only the right Mauthner fires, the left axon cap has reduced Kv1.1 and Cx35 localization (n=8/8 larvae with less Kv1.1 at left versus right cap; Figure 4C). Similarly, mutant larvae that are 100% right-based have diminished localization to the right axon cap (n=10/11 larvae with less Kv1.1 at right versus left caps; Figure 4D). We interpret from this data that spiral fiber axon misguidance in *celsr3* mutants stochastically leads to asymmetric input to the left and right Mauthner cells such that the Mauthner with the most spiral fiber input is preferentially activated, resulting in escapes only initiated in one direction. Thus, *celsr3* is required for proper spiral fiber targeting and maintaining balance within the startle circuit that enables animals to appropriately adjust their response to the spatial origin of startling stimuli.

### *celsr2* partially compensates for loss of *celsr3* in Mauthner development

In zebrafish, the Celsr gene family includes *celsr1a*, *1b*, *celsr2*, and *celsr3*, which have overlapping and unique expression patterns during nervous system development^34^. Previous studies have shown that Celsr3 and Celsr2 have partially overlapping roles in some neurodevelopmental contexts^22,53^. We hypothesized that the variable Mauthner phenotypes in *celsr3* mutants could be due to compensation by *celsr2*. *celsr2* mutants have do not have reduced Mauthner cell numbers at 5 dpf (Figure 5A-C,G), suggesting this gene is not required for Mauthner development. Similarly, *celsr2* mutants do not exhibit reduced escape frequencies or biased escapes (Figure 5H-J). However, loss of a single copy of *celsr2* in homozygous *celsr3* mutants strongly enhances the Mauthner cell loss phenotype (Figure 4G), and in >90% of *celsr3;celsr2* double mutant larvae, both Mauthner cells are absent at 5 dpf (Figure 5D-G). Spiral fiber neurons, however, are still present in *celsr3;celsr2* double mutants (Figure 5D-F). Additionally, we find that *celsr3;celsr2* mutants display a marked reduction of acoustic stimulus-induced responses (Figure 5H) and escapes (Figure 5I). This reduction is likely not due to the absence of Mauthners, as wild type larvae with bilateral Mauthner ablations perform Mauthner-independent, Mauthner-homolog mediated escapes with longer latencies than Mauthner-dependent escapes^54^. The occasional fast (≤17 ms latency) escapes we observe in *celsr3;celsr2* mutants lacking both Mauthners have a longer latency than those from *celsr3;celsr2* mutants with single Mauthners (Figure 5K). This indicates that Mauthner-homolog mediated escapes are at least partially intact in *celsr3;celsr2* mutants in lacking both Mauthners, and single Mauthner cells in *celsr3;celsr2* mutants are functional, as the escape latency in these larvae is within the range of Mauthner-dependent escapes. Combined, these data indicate that while *celsr2* is dispensable for normal Mauthner development, it can partially compensate for loss of *celsr3*.

**Figure 5:**
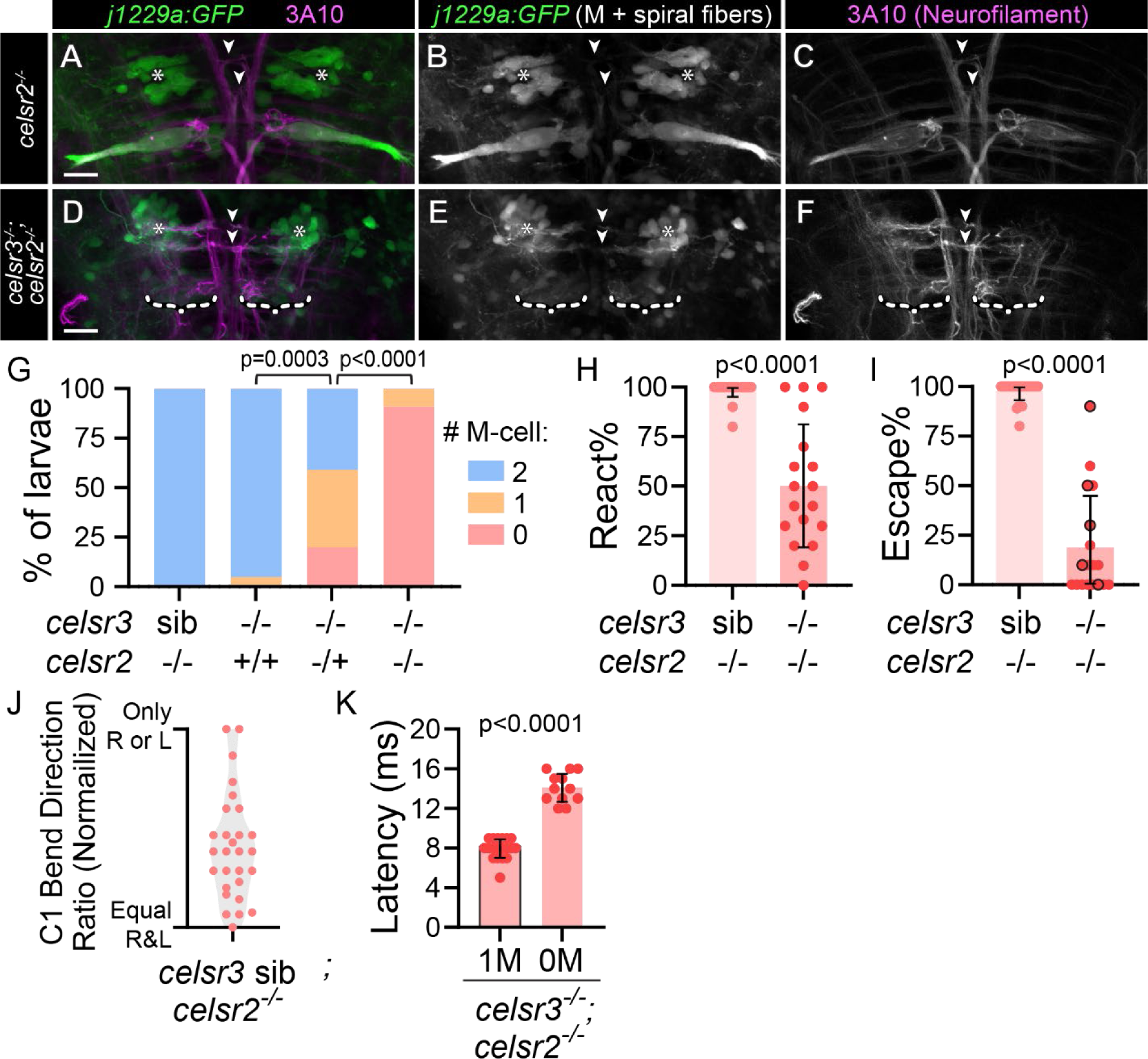
*celsr2* compensates for loss of *celsr3* during Mauthner development. A-F) *j1229a:GFP* transgene expression (A,B,D,E) labels Mauthner (A,B) and spiral fiber (asterisks; A,B,D,E) neurons in *celsr2^-/-^* (A-C) and *celsr3^-/-^;celsr2^-/-^* (D-F) larvae. α-3A10 (A,C,D,F) labels spiral fiber axons at midline crossing (white arrowheads) and Mauthner axons (A,C). White dashed brackets (D-F) indicate positions where Mauthner neurons are absent. All images and quantifications in figure are of 5 dpf larvae. Scale bar=20μM, Z-projection. G) Mauthner cell number in *celsr3* sib;*celsr2^-/-^* (100% 2 M-Cells; n=44), *celsr3^-/-^;celsr2^+/+^* (5% 1 M-cell, 95% 2 M-cells; n=20), *celsr3^-/-^;celsr2^+/-^* (20% 0 M-cell, 39% 1 M-cell, 41% 2 M-cells; n=41), and *celsr3^-/-^;celsr2^-/-^* (91% 0 M-cell, 9% 1 M-cell; n=22). p-values calculated with Chi-square tests. H) Percentage of twenty acoustic stimuli that elicited any response for *celsr3* sib;*celsr2^-/-^* (n=28) and *celsr3^-/-^;celsr2^-/-^* (n=17) larvae. I) For larvae from (H), percentage of twenty acoustic stimuli that elicited a fast escape (≤17 ms). Points outlined in black indicate *celsr3^-/-^;celsr2^-/-^* larvae with one Mauthner (5/17), while the remaining *celsr3^-/-^;celsr2^-/-^* larvae had zero Mauthners. J) For *celsr3* sib;*celsr2^-/-^* larvae from (I) that performed at least six escapes (100% of larvae), escape C1-bend ratio, normalized so all fish turning only one direction, left or right, appear at the same y position. K) Latency (time to first head movement) for individual fast escapes in *celsr3^-/-^*;*celsr2^-/-^* larvae with one (1M) or zero (0M) Mauthners. n=19 responses 1M, 12 responses 0M.

### Celsr3 is dispensable for facial branchiomotor neuron migration

Our findings that Celsr3 is required for development of two distinct neuronal populations, Mauthners and spiral fibers, prompted us to investigate Celsr3’s role in hindbrain neurons outside the startle circuit. Other PCP proteins, including Celsr2, are required in the hindbrains of both zebrafish^19^ and mice^22^ for facial branchiomotor neuron (FBMN) migration. The cell bodies of *islet-1/isl1+* FBMNs originate in hindbrain rhombomere 4 (r4), where Mauthner soma reside. By 48 hours post-fertilization (hpf), FBMNs have migrated to rhombomere 6 (r6) (Figure 6A)^55,56^. In zebrafish *celsr2* mutants, FBMNs fail to migrate and remain in r5 at 48 hpf^19^ (Figure 6B,E). In contrast, we find that by 48 hpf, FBMNs in *celsr3* mutants have appropriately migrated to r6 (Figure 6C,E). In addition, we fail to detect differences in the severity of migration deficits between *celsr2* single mutants and *celsr3;celsr2* double mutants (Figure 6D,E). Importantly, the lack of detectable differences between *celsr2* mutants and *celsr3;celsr2* mutants is unlikely to result from a ceiling effect in severity of migration as *celsr2* mutant larvae injected with a morpholino against *celsr1a/b* have a more severe migration phenotype than *celsr2* mutants alone^19^. Thus, while we cannot exclude a role for *celsr3* in FBMNs at later stages in development, as reported in mouse^22^, our data suggests that in zebrafish, *celsr3* is dispensable for FBMN migration.

**Figure 6:**
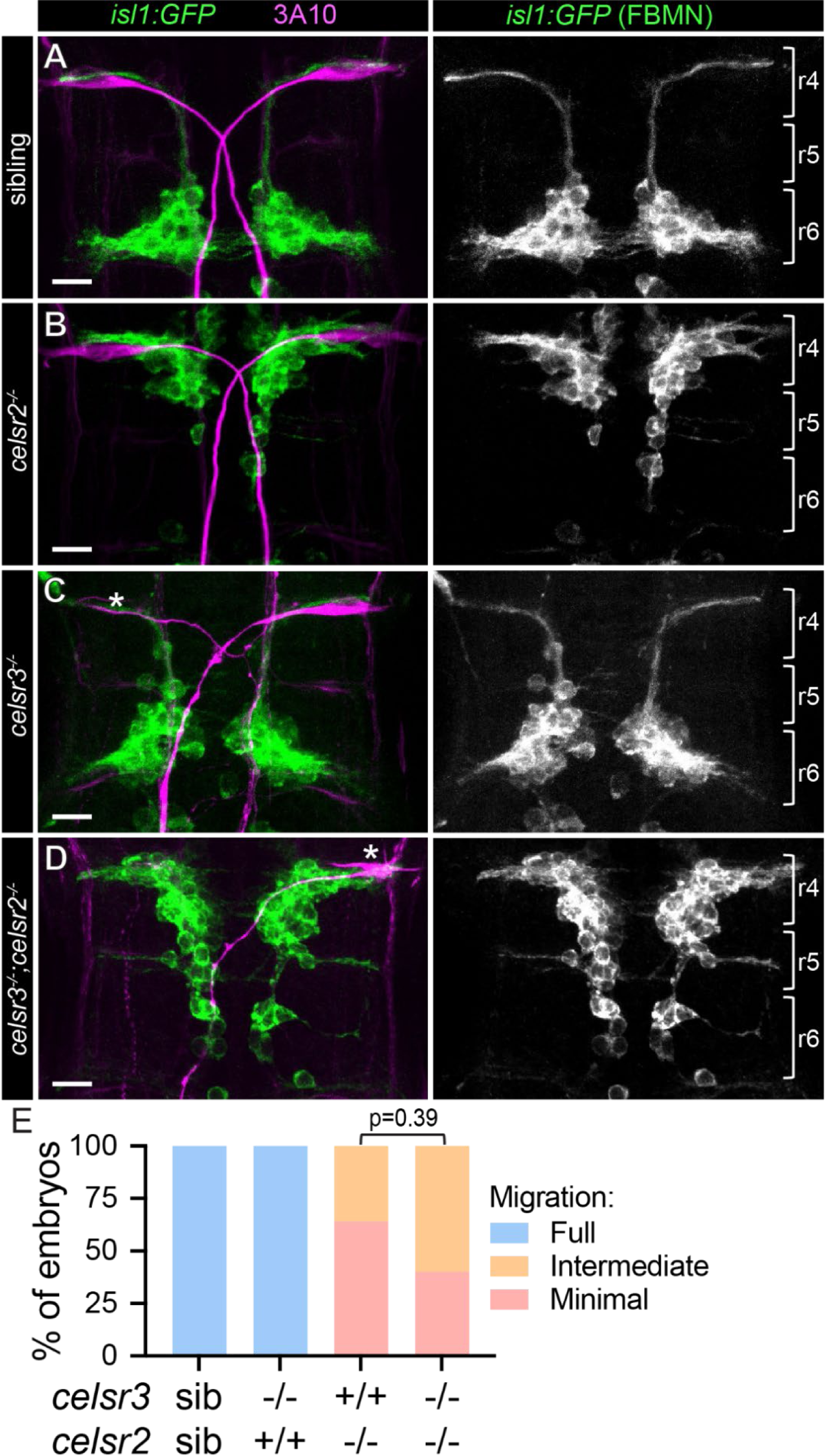
FBMN migration is independent of *celsr3*. A-D) *isl1:GFP* transgene expression of cytosolic GFP in 48 hours post-fertilization (hpf) facial branchiomotor neurons (FBMNs) in sibling (A), *celsr2^-/-^* (B), *celsr3^-/-^* (C), and *celsr3^-/-^;celsr2^-/-^* (D) embryos. α-3A10 labels Mauthner somas and axons (left panels). Approximate rhombomere boundaries (r4, r5, r6) are indicated with brackets (right panels). Asterisks in (C,D) denote presumptive dying Mauthners based on soma morphology. Scale bars=20 μM, Z-projection. E) Percent of larvae with full (e.g. A,C), intermediate (e.g. D), and minimal (e.g. B) FBMN migration. n=75 sib, 32 *celsr3^-/-^*, 11 *celsr2^-/-^*, 10 *celsr3^-/-^;celsr2^-/-^*. p-value calculated with Fisher’s exact test.

### Regional disruption of Celsr/Fzd signaling reveals a direct role for Celsr3 in spiral fiber axon guidance

Our results reveal that Celsr3 is required for both Mauthner development and spiral fiber axon guidance. One possible mechanism is that spiral fiber axon misguidance in mutants is a downstream effect of Mauthner cell developmental defects, rather than a direct role for Celsr3 in spiral fiber axon guidance. To distinguish between an indirect and direct role for Celsr3 in spiral fiber axon guidance, we sought to disrupt Celsr signaling specifically in rhombomere four (r4), where the Mauthner cell develops, and not r3, where spiral fiber soma reside. To achieve this, we utilized the Celsr3 binding partner, Frizzled 3a (Fzd3a)^23,57^, and regionally expressed a C-terminally truncated Fzd3a (Fzd3aΔC) that acts as a dominant negative^19,21^. Zebrafish have two *fzd3* genes, *fzd3a* and *fzd3b*, and *fzd3a* is the predominantly expressed homolog during embryogenesis^58^. *fzd3a* functions in a number of neurodevelopmental processes^19,21,59–61^, so we first asked if *fzd3a* is involved in startle circuit development. Similar to *celsr2* mutants, *fzd3a* single mutant larvae display normal startle behavior and Mauthner cell number (Figure 7A-C). In contrast, *celsr3;fzd3a* mutant larvae display similar, although milder, phenotypes compared to those we observe in *celsr3;celsr2* mutant larvae, with ∼69% of *celsr3;fzd3a* mutants lacking one or both Mauthner cells (Figure 7A-C). These data indicate Celsrs and Fzds work together to promote Mauthner development, and we predict disruption of Fzd3a function through expression of Fzd3aΔC will affect this Celsr/Fzd pathway.

**Figure 7:**
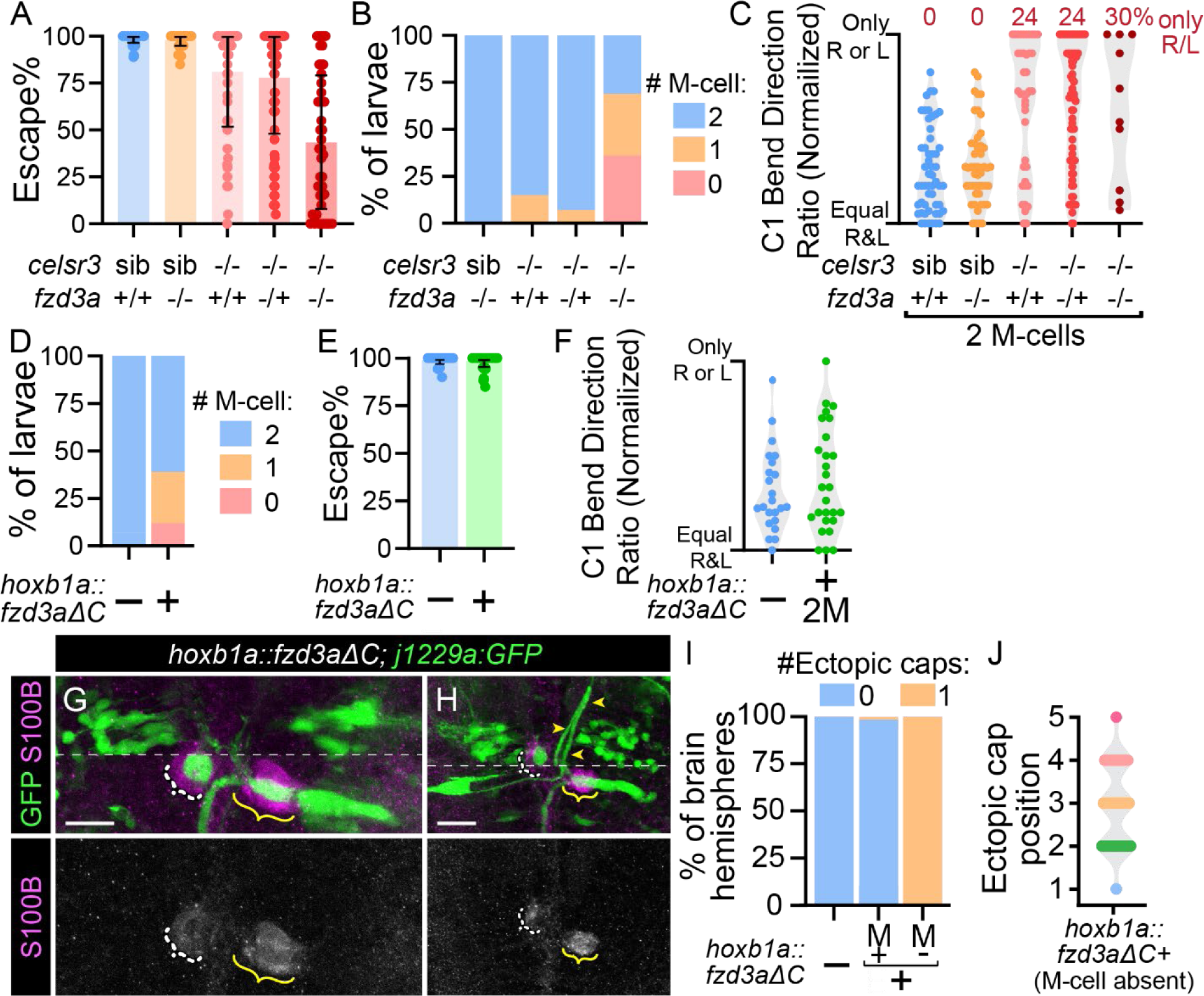
Spiral fiber axon guidance is partially independent from Mauthner development. A) Quantification of percent of twenty acoustic stimuli that resulted in a fast escape (≤17 ms). All images and quantifications in figure are of 5 dpf larvae. n=55 *celsr3* siblings, 51 *celsr3* sib;*fzd3a^-/-^*, 56 *celsr3^-/-^*, 92 *celsr3^-/-^;fzd3a^-/+^*, 45 *celsr3^-/-^;fzd3a^-/-^*. B) Mauthner cell number in larvae from (A). n=53 *celsr3* sib;*fzd3a^-/-^* (100% 2 M-cells), 55 *celsr3^-/-^* (15% 1 M-cell, 85% 2 M-cells), 86 *celsr3^-/-^;fzd3a^+/-^* (7% 1 M-cell, 93% 2 M-cells), 45 *celsr3^-/-^;fzd3a^-/-^* (36% 0 M-cells, 33% 1 M-cell, 31% 2 M-cells). C) Escape C1-bend ratio for larvae from (A) with two Mauthners (B) that performed at least six fast escapes in response to 20 acoustic stimuli, normalized so all fish turning only one direction, left or right, appear at the same y position. n=53 *celsr3* siblings, 48 *celsr3* sib;*fzd3a^-/-^*, 41 *celsr3^-/-^*, 71 *celsr3^-/-^;fzd3a^-/+^*, 10 *celsr3^-/-^;fzd3a^-/-^*. D) Mauthner cell number in control (no *hoxb1a:Gal4, UAS:fzd3aΔC/hoxb1a::fzd3aΔC* transgenes) and *hoxb1a::fzd3aΔC* larvae. n=28 control (100% 2 M-cells), 52 *hoxb1a::fzd3aΔC* (12% 0 M-cells, 27% 1 M-cell, 62% 2 M-cells). E) Quantification of percent of twenty acoustic stimuli that resulted in a fast escape (≤17 ms). n=30 control, 65 *hoxb1a::fzd3aΔC*. F) Escape C1-bend ratio for larvae from (E) with two Mauthners that performed at least six fast escapes in response to 20 acoustic stimuli, normalized so all fish turning only one direction, left or right, appear at the same y position. n=23 control, 27 *hoxb1a::fzd3aΔC*. G,H) *j1229a:GFP* labels Mauthner cells and spiral fiber neurons in *hoxb1a::fzd3aΔC* larvae while α-S100B labels glia. Yellow brackets indicate correctly positioned GFP+ spiral fiber axons at the Mauthner AIS, while white dashed brackets indicate displaced spiral fiber axon caps, due to either an absent Mauthner (G) or mispositioned Mauthner AIS (H). Yellow arrowheads in (H) denote aberrant path of left Mauthner axon, which continues posteriorly out of the Z-range displayed. Scale bars=20 μM, Z-projection (ventral anterior range and dorsal posterior range for GFP, for clarity; division marked by dotted line). I) Quantification of “ectopic caps,” as in Figure 3O,P. In brain hemispheres with no Mauthners, any large Kv1.1 accumulations were considered “ectopic” since no endogenous Mauthner AIS was present. n=16 control hemispheres, 60 *hoxb1a::fzd3aΔC* hemispheres with Mauthners, 32 hemispheres without Mauthners. J) Quantification of anterior-posterior position of ectopic caps from (K), as in Figure 3Q,R. n=32 *hoxb1a::fzd3aΔC* ectopic caps in hemispheres without Mauthners.

We next used *Tg(hoxb1a:Gal4)*^62^ to drive *UAS:fzd3aΔC* expression throughout r4 from ∼10-48 hpf^21^ and assessed whether Mauthner and spiral fiber neurons were disrupted. By 5 dpf, 39% of *hoxb1a::fzd3aΔC* larvae are missing one or both Mauthner cells (Figure 7D,G). In *hoxb1a::fzd3aΔC* larvae with two Mauthners, we did not detect defects in escape frequency (Figure 7E) or turn bias (Figure 7F), suggesting that spiral fiber neurons are functional. In the majority of *hoxb1a::fzd3aΔC* brain hemisphere with a Mauthner, Kv1.1+ ectopic caps were absent (n=59/60, quantified in Figure 7I), suggesting spiral fiber axons were not dramatically misguided as in *celsr3* mutants. In the rare *hoxb1a::fzd3aΔC* larvae with an ectopic or misplaced axon cap (n=4/60), the Mauthner soma was morphologically abnormal and lateral to its correct position (Figure 7H). When the Mauthner soma is incorrectly positioned and the spiral fiber axon cap is displaced, glia associate with the spiral fiber axon cap and not the Mauthner AIS (Figure 7G). We do not observe ectopically localized glia in *hoxb1a::fzd3aΔC* larvae (n=10). Additionally, in *hoxb1a::fzd3aΔC* brain hemispheres where the Mauthner was absent, a single “ectopic” cap was observed (Figure 7I,J), in contrast to *celsr3* mutants lacking a Mauthner cell (Figure 3P). This ectopic cap also forms closer to the correct position in *hoxb1a::fzd3aΔC* larvae (Figure 7J) than in *celsr3* mutant larvae (Figure 3R) and is associated with glia (Figure 7H). Together, these data indicate that spiral fiber defects present in *celsr3* mutants are unlikely a mere consequence of Mauthner cell defects. Instead, we conclude that *celsr3* is directly required for spiral fiber axon guidance.

### The PCP pathway promotes Mauthner anterior-posterior axon guidance

To identify whether the Celsr/Fzd pathway is required for the initial development or subsequent maintenance of Mauthner cells, we examined *hoxb1a::fzd3aΔC* embryos at 30 hpf. In age matched wild type animals, Mauthner cells have differentiated and their axons have extended across the midline and grown posteriorly into the spinal cord^63^. At 5 dpf, 39% of *hoxb1a::fzd3aΔC* larvae are missing one or both Mauthner cells (Figure 7D), while at 30 hpf, all *hoxb1a::fzd3aΔC* embryos have two Mauthner cells (Figure 8A-C). However, many Mauthner axons in these embryos have grown incorrectly in the anterior direction (Figure 8A-C). Since all fish have two Mauthners at 30 hpf, but only 39% still have two Mauthners at 5 dpf, we predict some Mauthner cells die between 30 hpf and 5 dpf due to axon guidance defects. We hypothesize that Mauthner cells that project axons anteriorly and fail to correct likely die, as we do not observe Mauthner axons growing solely anteriorly at 5 dpf. Consistent with this hypothesis, the ratio of 30 hpf embryos with two correct posteriorly growing axons versus one correct posterior and one incorrect anterior versus two incorrectly growing anterior is similar to the ratios of two versus one versus zero Mauthners in 5 dpf larvae (Figure 7D, 8C). We conclude that expression of Fzd3aΔC in r4 does not disrupt Mauthner specification but instead affects the process of axon growth and guidance, suggesting Celsr/Fzd signaling is required for Mauthner axon guidance.

**Figure 8:**
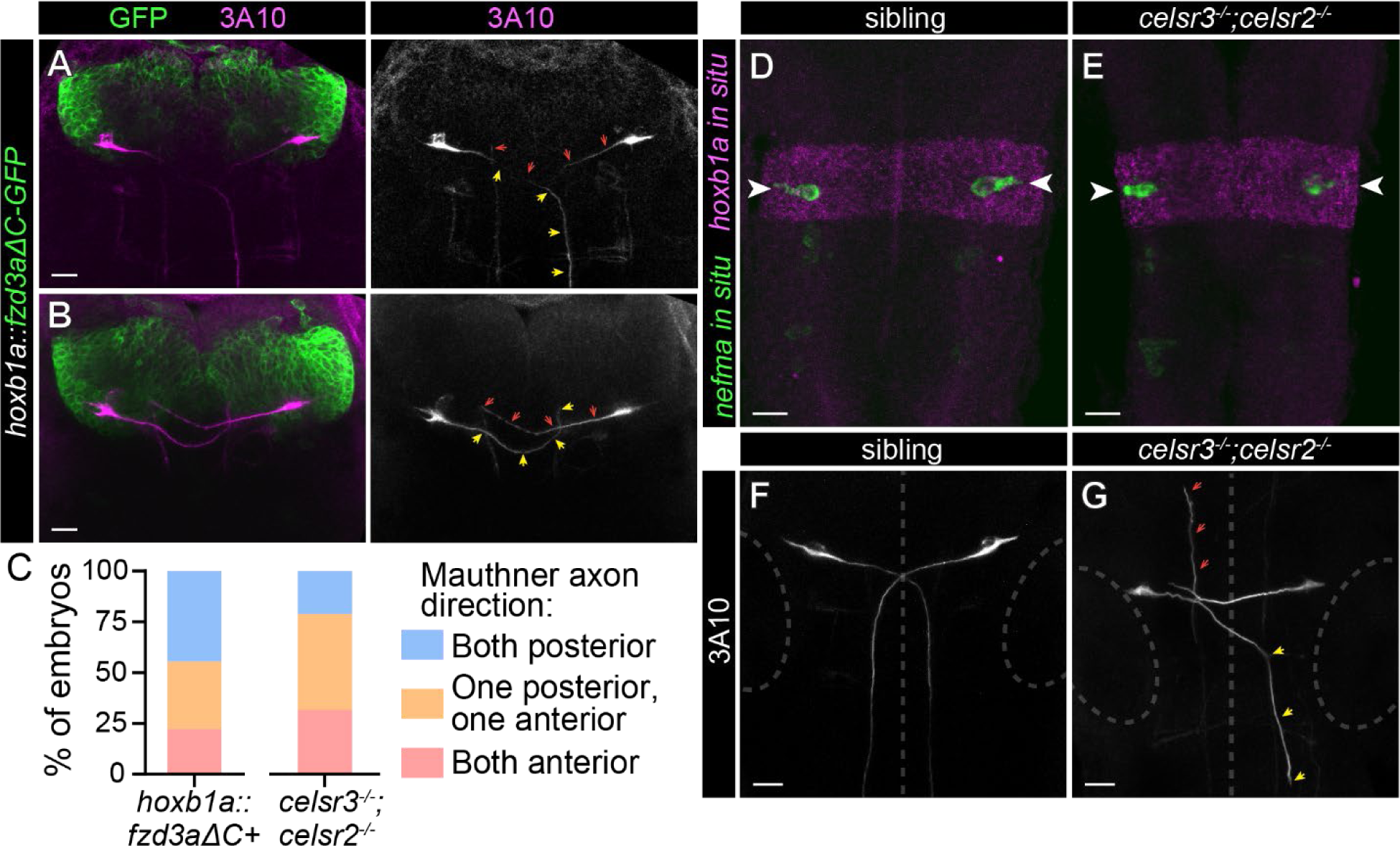
The PCP pathway directs A-P axon guidance of Mauthner cells. A-B) *hoxb1a::fzd3aΔC-GFP* transgene expression in hindbrain rhombomere 4. All images and quantification in figure are from 30 hpf embryos. α-3A10 labels Mauthner somas and axons. Yellow arrowheads indicate axons from left Mauthners and red arrows indicate axons from right Mauthners. In (A), the left axon grows correctly posteriorly while the right axon grows incorrectly towards the anterior, and in (B), both axons grow incorrectly towards the anterior. Scale bars=20 μM, Z-projection. C) Quantification of Mauthner axon phenotypes in *hoxb1a::fzd3aΔC* positive (n=18) and *celsr3^-/-^;celsr2^-/-^* (n=19) embryos. D,E) Sibling (D) and *celsr3^-/-^;celsr2^-/-^* (E) embryos with *hoxb1a in situ* (magenta) labeling hindbrain rhombomere 4 and *nefma in situ* (green) labeling reticulospinal neurons, including Mauthner cells (arrowheads). Scale bars=20 μM, Z-projection. F,G) Mauthner soma and axons are labeled with α-3A10 in siblings (F) and *celsr3^-/-^;celsr2^-/-^* (G) embryos. Yellow arrowheads indicate axon from left Mauthner growing correctly towards the spinal cord but displaced laterally. Red arrowheads indicate axon from right Mauthner growing incorrectly towards the anterior. Dotted lines indicate midline and encircle developing ears, for reference. Scale bars=20 μM, Z-projection.

Finally, we asked whether the Mauthner defects in *celsr3;celsr2* mutants were similarly correlated with axon guidance defects in early development. At 30 hpf, all *celsr3;celsr2* mutants have two Mauthner cells by gene expression (*in situ* for *neurofilament medium chain a*/*nefma* and *hoxb1a*, Figure 8D,E), indicating proper specification, and α-3A10 labeling (Figure 8F,G). While Mauthners are present and all Mauthner axons crossed the midline, many axons turned anteriorly rather than posteriorly (Figure 8C,G). Unlike *hoxb1a::fzd3aΔC*, in *celsr3;celsr2* mutants, more Mauthner cells are absent at 5 dpf (Figure 5G) than predicted if only Mauthner cells with anteriorly growing axons at 30 hpf die (Figure 8C). This suggests Celsrs promote additional developmental processes, such as progressive axon growth (Figure 2H), that ensure Mauthner cell survival. We conclude that the PCP pathway, particularly Celsr3 and Celsr2, promotes proper anterior-posterior Mauthner axon guidance.

## Discussion

Neural circuit assembly during development depends on numerous interacting molecular pathways driving axon targeting and connection of synaptic partners. Whether synapsing pairs of neurons within the same circuit utilize shared pathways to ensure connectivity, and what the consequences on circuit output are for altering but not abolishing these connections, is not fully understood. Here, we uncover a previously unappreciated role for the PCP pathway in the development and function of the acoustic startle hindbrain circuit. We find that Celsr3, along with Celsr2 and Fzd3a, are critical for axon growth and guidance of the Mauthner cells and spiral fiber neurons. Additionally, we demonstrate that disruption of spiral fiber guidance that leads to asymmetric input to the paired Mauthner cells has behavioral consequences leading to larvae unable to acutely determine escape direction, which is critical for escaping directional threats. Displaced spiral fibers axons also displace glia associated with spiral fiber-Mauthner synapses, suggesting a role for spiral fiber axons in glial recruitment to their target area, the axon cap. Finally, we discover divergent roles for Celsr3 and Celsr2 for hindbrain development with Celsr3 being the primary player in startle circuit development while Celsr2 plays a compensatory role, and Celsr2 being critical for FBMN migration while Celsr3 seems dispensable. Combined, these results provide compelling evidence for a critical role of the PCP pathway in assembly and connectivity of the acoustic startle circuit.

How do shared molecular pathways promote neural circuit connectivity? Cadherins, including those in the Celsr/Flamingo family, play various roles in axon growth and targeting. The clustered protocadherins have been especially well-studied in mammals for roles in self-avoidance and neurite tiling through cis and trans heterophilic and homophilic interactions, as well in synaptogenesis and neural connectivity through preventing inappropriate connections^64^. *Drosophila* Flamingo similarly plays roles in neurite tiling^26,27^ and preventing ectopic synapse formation^65^, as well as axonal fasciculation^66^, as does the *C. elegans* Flamingo homolog FMI-1^67^. In vertebrates, distinct roles in axon guidance and growth have been described for different Celsrs and different neural populations. In mammals, Celsr3 is required for anterior-posterior axis axon guidance in monoaminergic axons in the brainstem^68,69^ and commissural axons in the dorsal spinal cord^70^, likely through intercellular interactions between PCP proteins promoting growth cone turning^23,71,72^. Celsr3 has also been implicated in synapse stabilization^28^ and inter-growth cone interactions that promote axon organization and progressive axon growth^69^. In future studies, it will be interesting to determine whether Celsr3 homophilic interactions between Mauthner and spiral fibers neurons similarly promote their synaptic connectivity, especially as this connection offers the opportunity to investigate Celsrs in axo-axonic synapse formation.

In addition to divergent roles for different classes of cadherins, there are also distinct functions within the Celsr family. Here, we report that Celsr3 and Celsr2 are differentially required in two developmental processes in the hindbrain: startle circuit assembly and FBMN migration. This may reflect differences in developmental timing for these two processes, as Mauthner axon turning along the anterior-posterior axis occurs several hours before FBMN migration, or differential expression in different neuron populations, though *celsr3* and *celsr2* expression in the hindbrain largely overlap at this point in development^34^. Alternatively, these distinct requirements may reflect differential functions of the proteins. While Celsrs have similar protein structures^32,34^, differences do exist, particularly in the C-terminal intracellular domain. Though the function of these intracellular domains is largely unknown, C-terminally truncated Celsr2 in zebrafish is sequestered in the golgi, and expression of just the C-terminal domain with a membrane tag disrupts convergence/extension in embryos^73^. Additionally, recent work has demonstrated that mammalian CELSRs differentially engage G proteins, which may result in differential G protein-dependent signaling^74^. Future experiments in zebrafish could address whether the differential roles for Celsr3 and Celsr2 in hindbrain development can be attributed to differences in protein sequence.

Intracellular signaling in response to PCP pathway activity is thought to support growth cone turning by promoting cytoskeletal rearrangement^23^. The ligand that induces this PCP activity is not entirely clear, and may differ for different neuron populations, but Wnt^75,76^ and ephrinA^57^ proteins act in specific axon guidance processes. Both Wnt proteins, which interact with Fzd receptors, and Eph/Ephrin pairs, which can act as a co-receptors for Fzds, are expressed in distinct rhombomere and rhombomere boundaries in the zebrafish hindbrain^77,78^, making them potential upstream regulators for Celsr3-mediated Mauthner and spiral fiber axon guidance. Downstream of PCP receptor/ligand interactions, the intracellular signaling that leads to cytoskeletal rearrangements in growth cones likely overlaps with proteins involved in other PCP-dependent processes, including Disheveled proteins^71,72^. Formin proteins, which help regulate actin polymerization, are also implicated as downstream PCP regulators^79,80^. Interestingly, *formin2b*/*fmn2b* zebrafish morphants and crispants have reduced spiral fiber commissures and perform reduced short latency escapes^81^, similar to the phenotype of wild type fish with ablated spiral fibers^13^. This data suggests studies on spiral fiber axon development may provide insight on molecular and genetic interactions between *celsr3* and *fmn2b*, as well as additional proteins proposed to act during axon guidance. Additionally, the ability to tag proteins and visualize axonal dynamics *in vivo*, particularly in the uniquely large Mauthner cell, may offer an invaluable in vivo insight to localization of various PCP proteins during the process of growth cone turning and axon growth.

Neural circuit assembly is orchestrated by a myriad of genetic and molecular pathways. Simple and well-characterized sensorimotor circuits like the acoustic startle circuit offer an opportunity to untangle these complex molecular interactions and identify relationships between synapsing pairs of neurons and specific cellular pathways. In addition to identifying the first, to our knowledge, pathway regulating anterior-posterior guidance in Mauthner cells, our work also provides compelling evidence that the zebrafish hindbrain is a prime model for developmental studies on molecular mechanisms for PCP-directed circuit assembly.

## Methods

### Key Resources Table

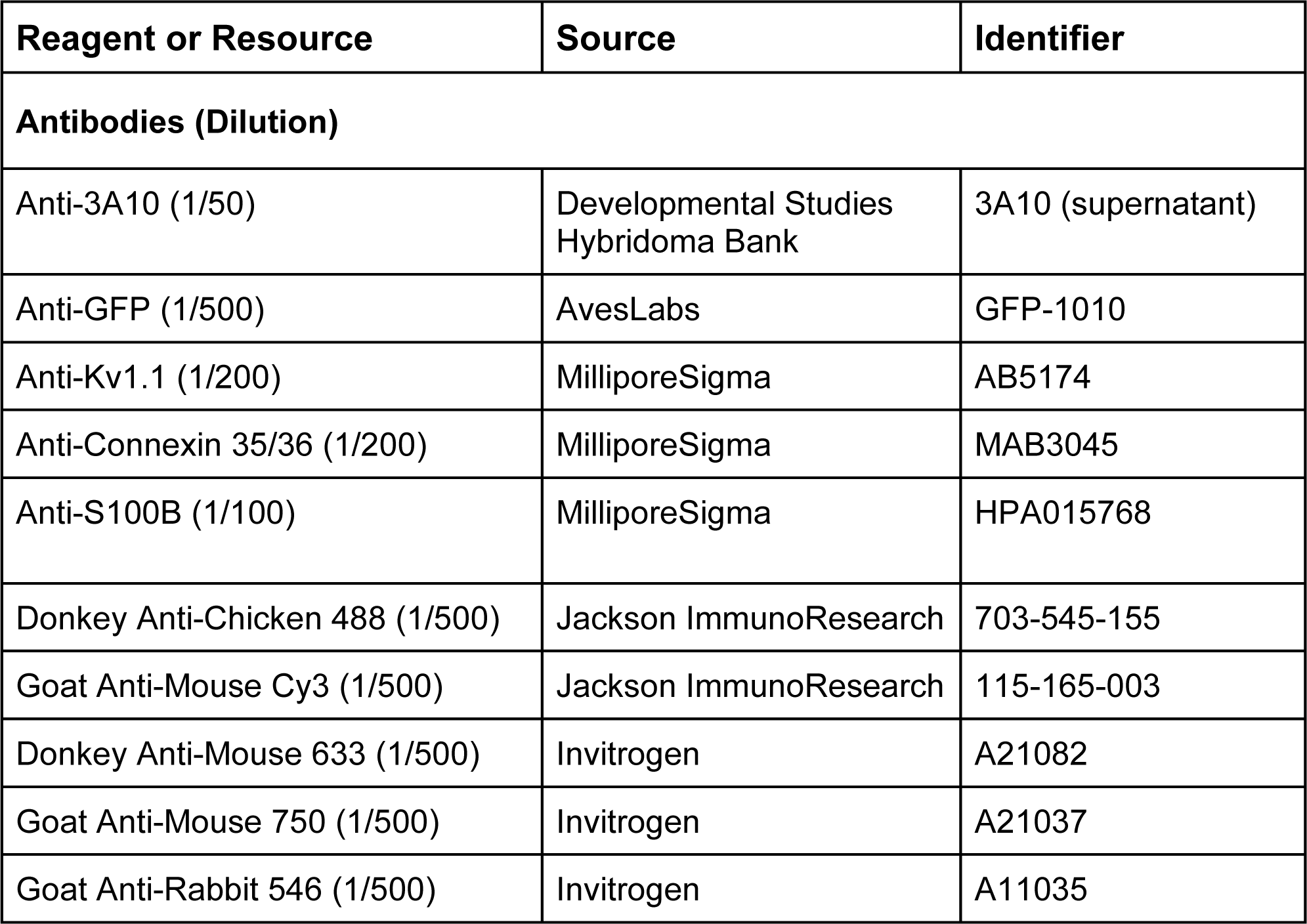
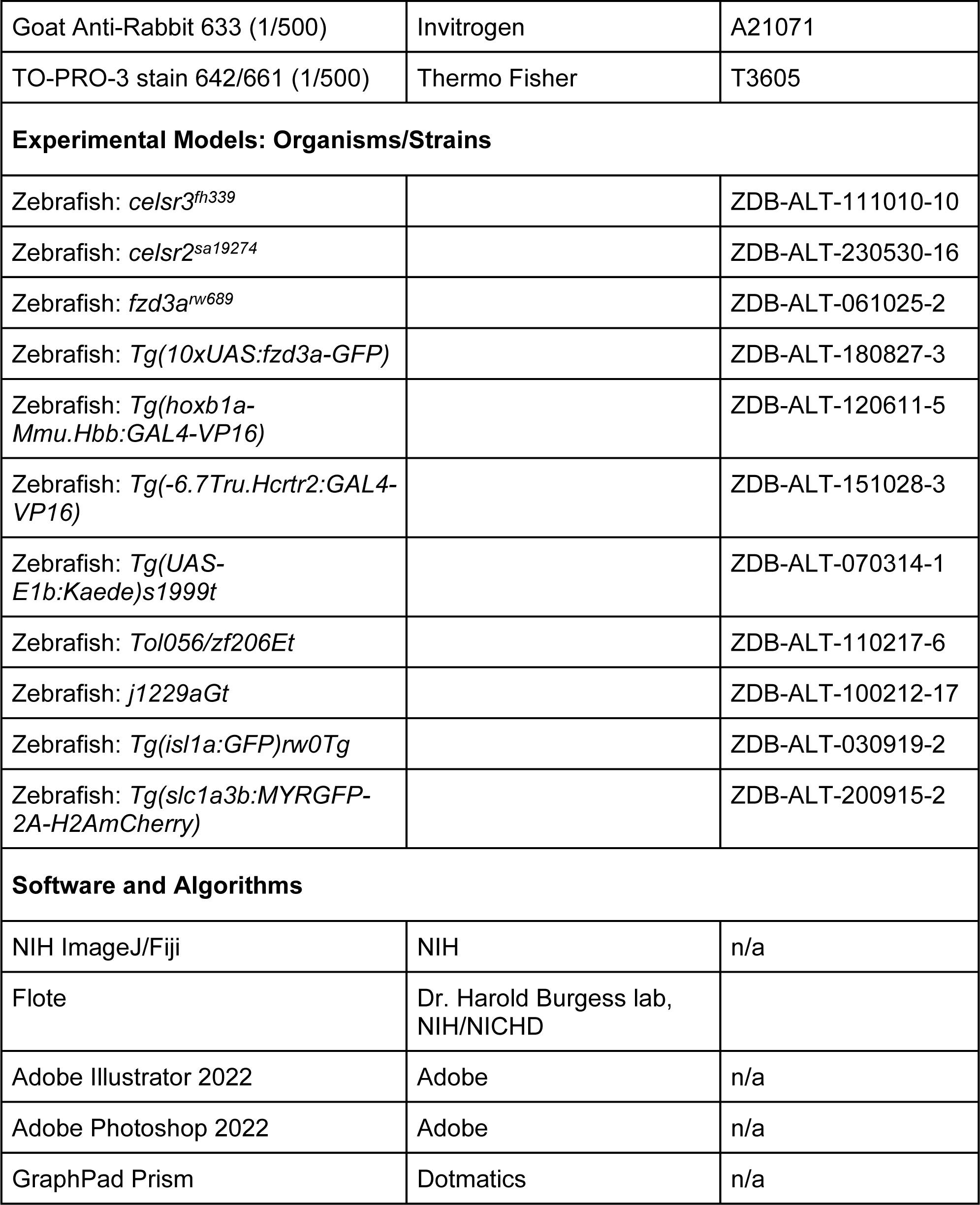
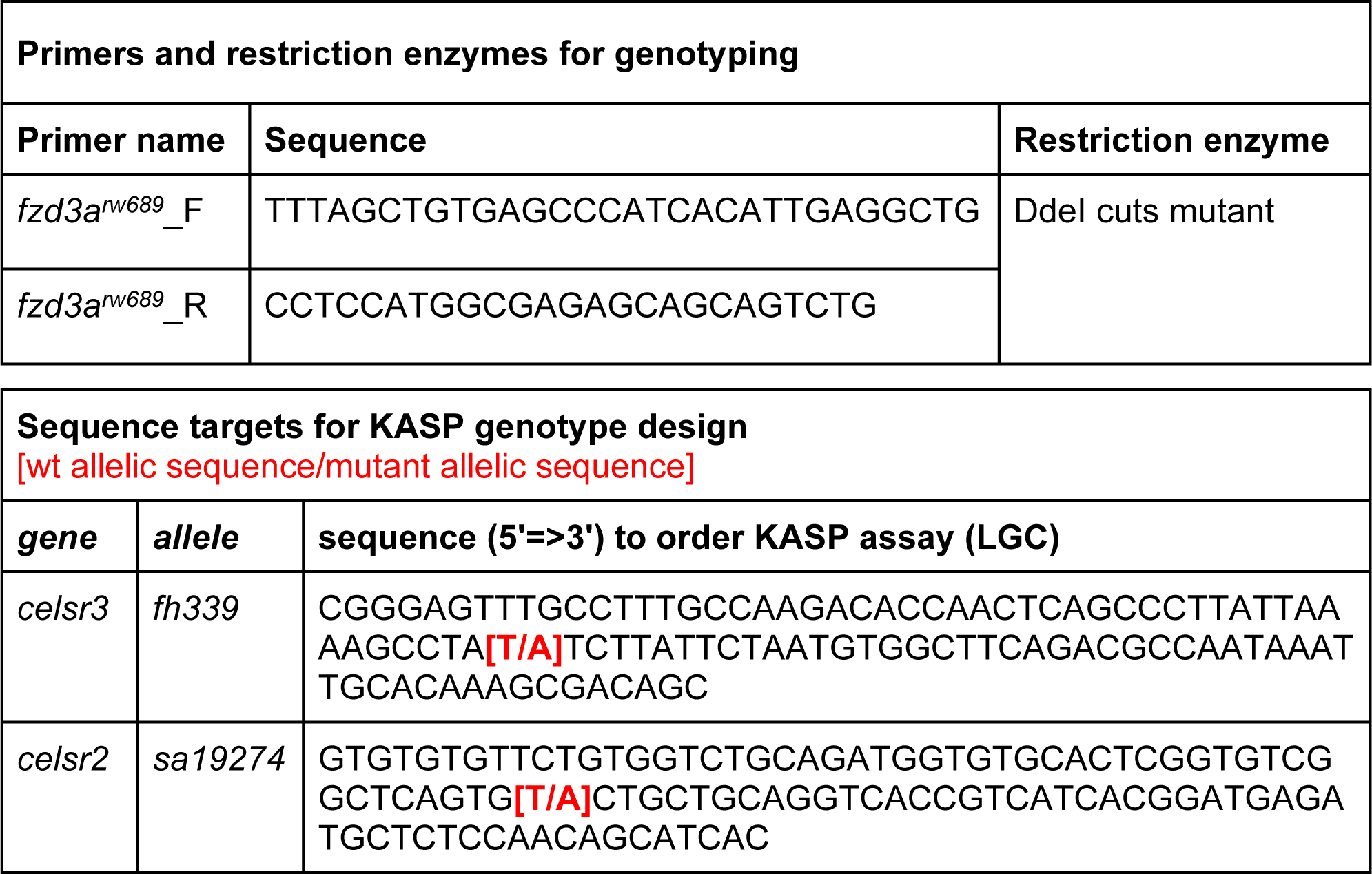

### Lead Contact and Materials Availability

Further information and requests for reagents and resources should be directed to and will be fulfilled by the Lead Contact, Michael Granato.

### Experimental Model and Subject Details

#### Zebrafish and Maintenance

All animal protocols were approved by the University of Pennsylvania Institutional Animal Care and Use Committee (IACUC). Embryos were raised at 29°C on a 14-h:10-h light:dark cycle in E3 media. Mutant and transgenic lines used in this study are listed in the Key Resources Table.

### Method Details

#### Behavior testing

Behavioral experiments were performed using 5 dpf larvae and analyzed using Flote software^15^. Larvae were tested in groups in a 6×6 laser cut grid. ∼25.9 dB acoustic stimuli were used for acoustic startle experiments, based on larval response rates and previous measurements of the speaker^39^. 20 stimuli were given with a 20 second interstimulus interval. Videos of acoustic startle response were acquired at 1000 frames per second (fps). Before Flote tracking and analysis, NIH ImageJ/Fiji was used to remove background from videos to prevent erroneous measurements from the outline of the wells. Larvae with >2 untracked responses due to tracking errors were excluded from further analysis. Escape% per Mauthner was calculated with the following formula: [Escape% for a left Mauthner]=[Escape%, larva] x [%Right turns, larva] and [Escape% for a right Mauthner]=[Escape%, larva] x [%Left turns, larva]. For example, for a larva with 80% escape response, and 60% right turns, we calculate a left Mauthner escape% of 48% and a right Mauthner escape% of 32%.

#### Immunohistochemistry

5 dpf larvae were fixed with either 2% trichloroacetic acid (TCA) for 3 hours at room temperature (Kv1.1 staining) or 4% paraformaldehyde (PFA) overnight at 4°C (all other staining) and stained as previously described^82^ (Martin et al, bio-protocol 2022). Before mounting in Vectashield, TCA-fixed brains and spines were peeled^83^, and PFA-fixed brains were dissected. Antibodies and dilutions used are listed in Key Resources Table.

#### Imaging and quantification

Mauthner soma numbers were quantified in live *Tol-056* larvae using an Olympus MVX10 fluorescent stereo microscope. Mauthner axon lengths and glia localization were quantified in live *Tol-056* larvae using an Olympus IX81 spinning disk confocal with a 40x lens. Fixed samples for figure images and quantification were acquired on a Zeiss LSM 880 or 980 laser scanning confocal with a 20x lens.

#### Genotyping

See Key Resources Table for details.

#### Statistics

Statistical analyses were performed using GraphPad Prism. P-values were calculated using non-parametric Mann-Whitney test, unless otherwise noted in figure legend. Bar graphs display mean and standard deviation.

## Acknowledgements

The authors would like to thank Drs. Cecelia Moens, Greg Walsh, Mary Mullins, and David Schoppik for zebrafish lines; the UPenn Cell & Developmental Biology Microscopy Core; and members of the Granato lab for feedback regarding the manuscript.

## Author Contributions

Conceptualization: J.H.M., M.F.N., M.G.

Methodology: J.H.M., M.F.N., E.A.O.

Investigation: J.H.M., M.F.N., E.A.O.

Formal analysis: J.H.M., M.F.N., Visualization: J.H.M., M.F.N.

Writing-original draft: J.H.M., M.F.N.

Writing-review and editing: J.H.M., M.F.N., E.A.O., M.G. Funding acquisition: M.G.

